# Virus-induced transgene- and tissue culture-free heritable genome editing in tomato

**DOI:** 10.1101/2025.11.08.687402

**Authors:** Ye Liu, Trevor Weiss, Jinhee Lee, Jessica Powell, Shi Ying Charlize Choo, Elnaz Roshannai, Maris Kamalu, Jasmine Amerasekera, Suhua Feng, Steven E. Jacobsen

## Abstract

Genome editing has emerged as a powerful tool for genome manipulation and trait improvement in crops. However, most commonly used approaches rely on tissue culture and transgenic materials, which are time-consuming, labor-intensive, and often strongly genotype-dependent. Here, we developed a tobacco rattle virus (TRV)–based system to deliver the compact ISYmu1 TnpB endonuclease, coupled with *in planta* shoot regeneration, to achieve somatic and heritable genome editing across different tomato cultivars without tissue culture. By targeting *SlPDS*, we successfully generated virus-free homozygous mutant progeny in a single generation. Furthermore, we extended this system to the functional analysis of the previously uncharacterized *SlDA1* locus, revealing its involvement in organ size regulation, and recovered transgene-free *SlDA1* mutants displaying enlarged fruits. Given the wide host range of TRV, our system should be broadly applicable for rapid, non-transgenic and less genotype-dependent heritable genome editing, thereby advancing both functional genomics and crop improvement.

**Significance:** Efficient genome editing without the need for transgenesis or tissue culture remains a major challenge in crop breeding. Here, we establish a simple single-step system for transgene- and tissue culture-free genome editing in tomato based on Tobacco Rattle Virus-mediated delivery of the compact RNA-guided TnpB enzyme ISYmuI and its guide RNA. This strategy enabled somatic editing of *de novo* shoots and heritable transmission of targeted mutations to the next generation. Notably, editing an agronomically relevant gene produced tomato plants with larger fruits, highlighting the potential of this system for crop improvement.

## Introduction

The emergence of genome editing technology offers tremendous potential for accelerating crop improvement(1, 2). However, despite the rapid adoption of genome editing in many major crops, two significant bottlenecks remain: efficient delivery of genome-editing reagents and the efficient generation of transgene-free plants. Conventional plant genome editing often relies on the use of tissue culture and plant transformation, which typically involves introducing the editing construct into plants through *Agrobacterium tumefaciens*-mediated transformation, selecting transgenic plants carrying the desired mutation, and subsequently removing the transgenic components by crossing(3, 4). This method is time- and labor-intensive and is limited to specific crop species and genotypes, thereby hindering the application of genome editing in crop plants.

To overcome these barriers, several plant viruses have been engineered to deliver single-guide RNAs (gRNAs) directly to transgenic plants expressing SpCas9 for virus induced genome editing (VIGE) in both dicotyledonous and monocotyledonous crop species(5–9). For example, some RNA viral vectors, such as tobacco rattle virus (TRV) and barley stripe mosaic virus (BSMV) have been shown to deliver gRNA to meristematic and germline cells to achieve heritable editing, in part with the help of RNA mobility sequences(10–13). However, most plant viruses are excluded from meristematic tissues, limiting the access to the germline and thus reducing the potential to generate gene-edited progeny (14, 15). Moreover, due to the restricted cargo capacity of most viruses, the application of these methods still requires the additional steps of using tissue culture and transformation to obtain the overexpressing-Cas9 line as well as crossing to segregate away the transgenes after edits are obtained(16).

A recent strategy to circumvent these limitations was to engineer TRV to deliver the compact ISYmu1 TnpB nuclease and the gRNA, to enable transgene-free germline editing in *Arabidopsis*(17). It remains unknown, however, whether this approach can be applied to crop species. Alternatively, large-capacity vectors such as barley yellow striate mosaic virus (BYSMV) have been used to deliver Cas9 and gRNAs for heritable genome editing in wheat(18, 19). However, this approach faces limitations due to its complexity and dependence on insect transmission, making it difficult to adapt to other plant species. Thus, the development of simple, rapid, and broadly applicable methods for producing transgene-free, heritable genome-edited crops without the need for tissue culture remains an urgent priority.

In this study, we established a strategy for TRV-mediated delivery of the compact ISYmu1 TnpB endonuclease and gRNAs in tomato. Using this system, we first targeted the *SlPDS* gene to successfully generate somatically edited shoots, and also observed transmission of edited alleles to the next generation. We further applied this system to the previously uncharacterized *SlDA1* locus, and generated mutants with enlarged fruits. Together, these results indicate that our platform enables efficient somatic editing and stable transmission of targeted mutations in a crop, thereby providing a streamlined, one-step strategy for heritable, transgene-free genome editing in tomato that bypasses tissue culture.

## Results

### Viral Delivery of ISYmu1 for Somatic Genome Editing in Tomato

Given the ultracompact size of TnpBs, it represents an attractive nuclease for virus-induced genome editing, as viral vectors can deliver and express the entire genome-editing reagents(17), thereby enabling editing without the need for Cas9-overexpressing plants and eliminating the requirement for transgenesis and tissue culture in crops such as tomato. To this end, we engineered a TRV vector to express both ISYmu1 TnpB endonuclease and its guide RNA, fused to a truncated mutant FT sequence of tomato (mSIFT, with a mutation in the start codon) (Fig. 1A), which has been shown to enhance mobility of the gRNA between cells(20). We then generated a pTRV2-TnpB-gRNA^SlPDS^-mSlFT vector that expresses a gRNA targeting the tomato phytoene desaturase gene (*SlPDS*), which was delivered to a Red cherry-type cultivar via an *Agrobacterium tumefaciens* cotyledon infiltration method (Fig. 1B, S1A). Three weeks post-infiltration, DNA was isolated from young leaves and used as a template for PCR, followed by NGS analysis of PCR amplicons flanking the *SlPDS* target site. Of the 20 plants infiltrated, we observed three (15%) plants with somatic editing, ranging from 0.8%∼6.93% indels in systemically infected leaves (Fig. S1B). We collected seeds from #13 plant showing 6.93% somatic editing to screen for heritability of edited alleles, however, these somatic mutations failed to be transmitted to the progeny seedlings, which may be due a combination of low efficiencies of agrobacterium delivery, viral spread, the efficiency of TnpB activity, and/or instability of the engineered TRV RNA2. Collectively, these data indicated that TRV can deliver ISYmu1 and gRNA to induce somatic genome editing in tomato.

**Fig. 1.**
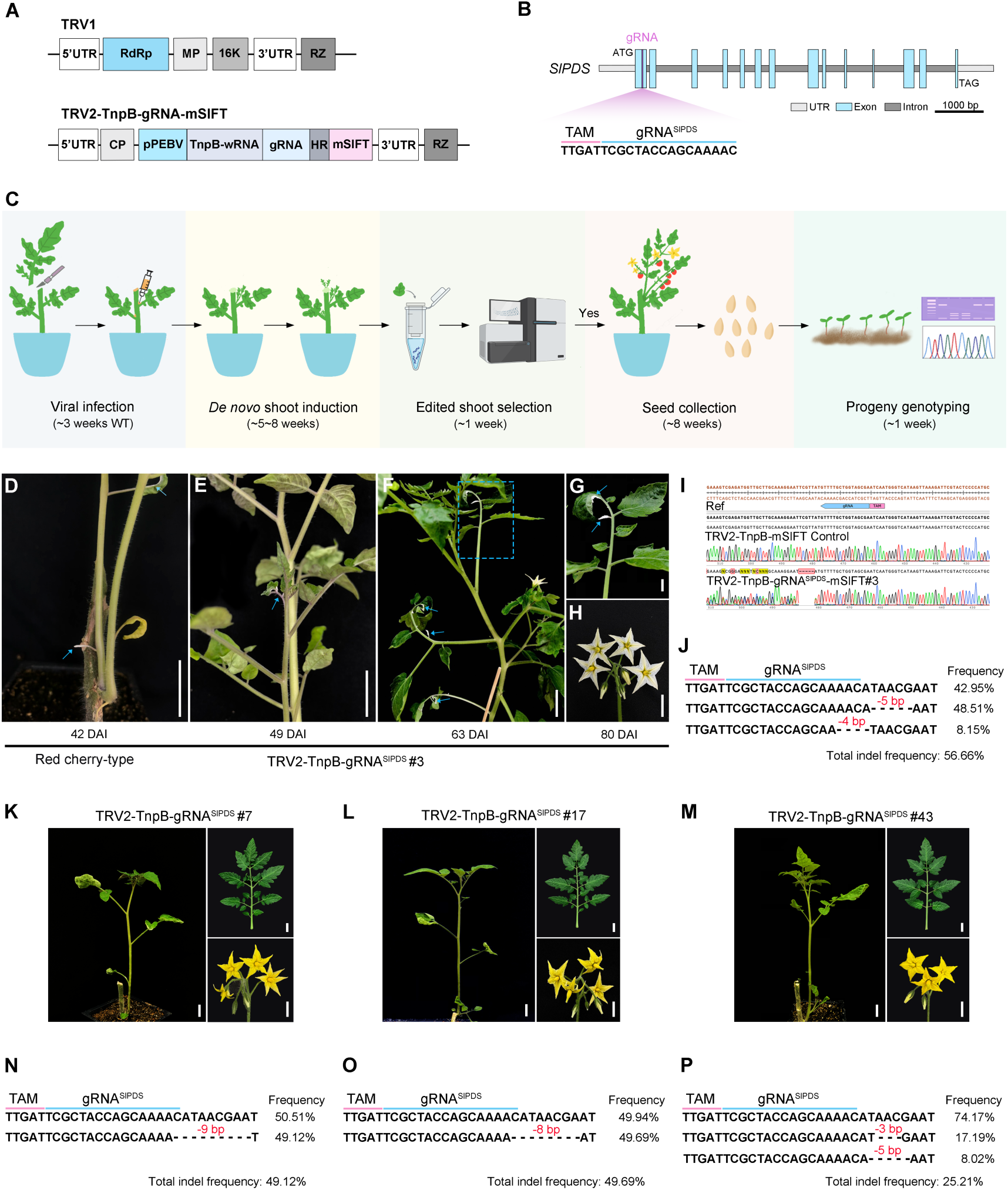
Virus-induced genome editing in *de novo shoots* of tomato using TRV-TnpB-gRNA^SlPDS^-mSlFT. (**A**) Schematic diagram of the TRV1 and TRV2 plasmids. The tomato mutated *FT* mobility element (mSlFT) was fused to the 3’ end of the gRNA in TRV2. RdRp, RNA-dependent RNA polymerase; MP, movement protein; CP, coat protein; pPEBV, *Pea Early Browning Virus* subgenomic promoter; HR, HDV ribozyme, mSlFT; mutated tomato *FT* mobility element. (**B**) Schematic representation of *SlPDS* genomic structure. The light blue box represents exons, light grey represents the UTR sequence, and dark grey represents introns. The TAM sequence and target sequence are highlighted by the pink and blue lines, respectively. (**C**) The procedure for TRV-TnpB-gRNA-mSlFT mediated genome editing system in tomato plants. Edited *de novo* shoots were screened and validated by Sanger and NGS amplicon sequencing. (**D-H**) Different growth stages of TRV-TnpB-gRNA^SlPDS^#3 plant displaying albino sectors in leaves and flowers. White leaflets in leaves are indicated by blue arrows, and the flowers exhibited white petals (**H**). **G** is a close-up view of **F**; DAI: Days After Injection. (Scale bars, 1 cm). (**I**) Sanger sequencing of the *SlPDS* gene target site in TRV-TnpB-mSlFT Control (top) and TRV-TnpB-gRNA^SlPDS^#3 (bottom) plants. (**J**) Indel frequency analysis of the *SlPDS* gene in the TRV-TnpB-gRNA^SlPDS^#3 shoot. Genomic DNA was extracted from green leaf tissue and subjected to amplicon sequencing using next-generation sequencing (NGS). The TAM sequence and target sequence are highlighted by the pink and blue lines, respectively. Deleted residues are marked by black dotted lines. The different indel types are shown on the left. The percentage of reads corresponding to each indel type is shown on the right. The total indel efficiency was calculated by the total number of indel reads divided by total number of reads. (**K-M**) Phenotype of the TRV-TnpB-gRNA^SlPDS^#7 (**K**), #17 (**L**) and #43 (**M**) shoots, respectively, showing no obvious albino sectors. Images include the whole plant (left), a leaf (top right) and flowers (bottom right). (Scale bars, 1 cm). (**N-P**) Indel frequency analysis of the *SlPDS* gene from the TRV-TnpB-gRNA^SlPDS^#7 (**N**), #17 (**O**) and #43 (**P**) shoots. The reads of sequences with frequencies >5% are shown as well as the total indel frequency.

### Induction of Genome Edited *De Novo* Shoots via Viral Delivery of ISYmu1 and gRNA^SlPDS^

Emerging *in planta* regeneration approaches offer the potential for more efficient induction of *de novo* shoots(21, 22). This inspired us to explore a heritable editing strategy in which genome editing was first induced in somatic cells, followed by the regeneration of *de novo* shoots from the edited cells. To achieve this, we removed the shoot apical meristem and all axillary shoots from Red cherry-type tomato plants, and then injected *Agrobacterium* encoding TRV-TnpB-gRNA^SlPDS^-mSlFT into all wounding sites of the main stem and axillary regions (Fig. 1C, S2 A-C). Five weeks after injection, we observed new shoots initiating from axillary regions, or the formation of callus-like tissues, from which additional shoots subsequently emerged, suggesting that *de novo* shoots arose from either direct or indirect regeneration, or both (S2 D, 2 E). A total of 49 plants were injected, from which 72 *de novo* shoots were initiated (Table S1). One of the *de novo* shoots (#3) showed full or partial white sectors on the plant similar to the known *SlPDS* mutant phenotype (Fig. 1D-G), suggesting biallelic mutations in the *SlPDS* gene. To confirm this, we first performed Sanger sequencing on three randomly selected green leaves of this shoot, which all revealed a 5-bp deletion as indicated by the presence of two sets of peaks in the sequencing traces at the target site (Fig. 1I). Further amplicon-sequencing (amp-seq) analysis showed that this plant exhibited 56.66% somatic editing efficiency in green leaves (Fig. 1J), consisting of 48.51% of the 5-bp deletion and another 8.15% with a 4-bp deletion. Interestingly, as the plant grew, the petals of all the flowers were white (Fig. 1H), but the fruit did not display a photobleaching phenotype. We also isolated genomic DNA from different organs to measure the mutation frequency. Amp-seq analysis showed that different tissues displayed distinct mutation patterns and editing efficiencies (S3A, 3B), indicating that TRV-TnpB-gRNA^SlPDS^#3 was chimeric and that genome editing was continued to occur late in the development of this shoot.

For other newly developed shoots without a photobleaching phenotype (Fig. 1K-M), we first performed Sanger sequencing as a preliminary screening. The results showed mixed peaks in TRV-TnpB-gRNA^SlPDS^#7, #17, #43 and #70 lines out of 71 shoots (S4A-F), suggesting genome editing had occurred. To quantify this editing, tissue samples from these lines were collected for amp-seq analysis, which revealed somatic mutation frequencies ranging from 25.21% to 49.69% (Fig. 1N-P, S5). Notably, #7 and #17 shoots appeared to carry a single predominant deletion with a frequency of roughly 50% (Fig. 1N, 1O), suggesting these lines were heterozygous for a mutant allele that likely arose during early development of the *de novo* shoot. These data demonstrate that TRV-TnpB-gRNA^SlPDS^-mSlFT can induce a high level of somatic edits in *de novo* shoots.

### Heritable Genome Editing via Viral Delivery of ISYmu1 and gRNA^SlPDS^

We next investigated whether targeted mutations could be transmitted to the next generation. A total of 96 seeds from four lines (#3, #7, #17 and #43) were sown on 1/2 MS plates. After one week, albino seedlings were observed in the progeny of shoots #3 and #17, suggesting biallelic mutations in the *SlPDS* gene (Fig. 2A, 2B, 2E and 2G). To characterize the genotypes of the seedlings, we performed Sanger sequencing. Analysis of seedlings from shoot #3 revealed 27 homozygous mutants (28%) with a 5-bp frame-shift deletion, 39 heterozygous mutants (41%) and 30 wild type (31%), consistent with a roughly 3:1 segregation ratio and confirming the parent plant was likely heterozygous for the 5-bp deletion (Fig. 2I). Similarly, the seedlings of shoot #17 showed 23 homozygotes for a 4-bp deletion (24%), 54 heterozygotes (51%) and 19 wild type (20%) (Fig. 2I), and shoot #7 produced 27% homozygotes (9-bp deletion) and 51% heterozygotes. The progeny of shoot #47, however, showed a lower transmission of mutations, with only 2% homozygotes and 24% heterozygotes for a 3-bp deletion. It is not surprising that we did not observe albino plants in the progeny of #7 and #43 plants (Fig. 2B, 2D, 2F, 2H), since these lines carried 9-bp or 3-bp deletions, respectively, which would be in-frame deletions. Together these data indicate that TRV delivery of ISYmu1 and gRNA fused to mSlFT is capable of inducing heritable editing in tomato.

**Fig. 2.**
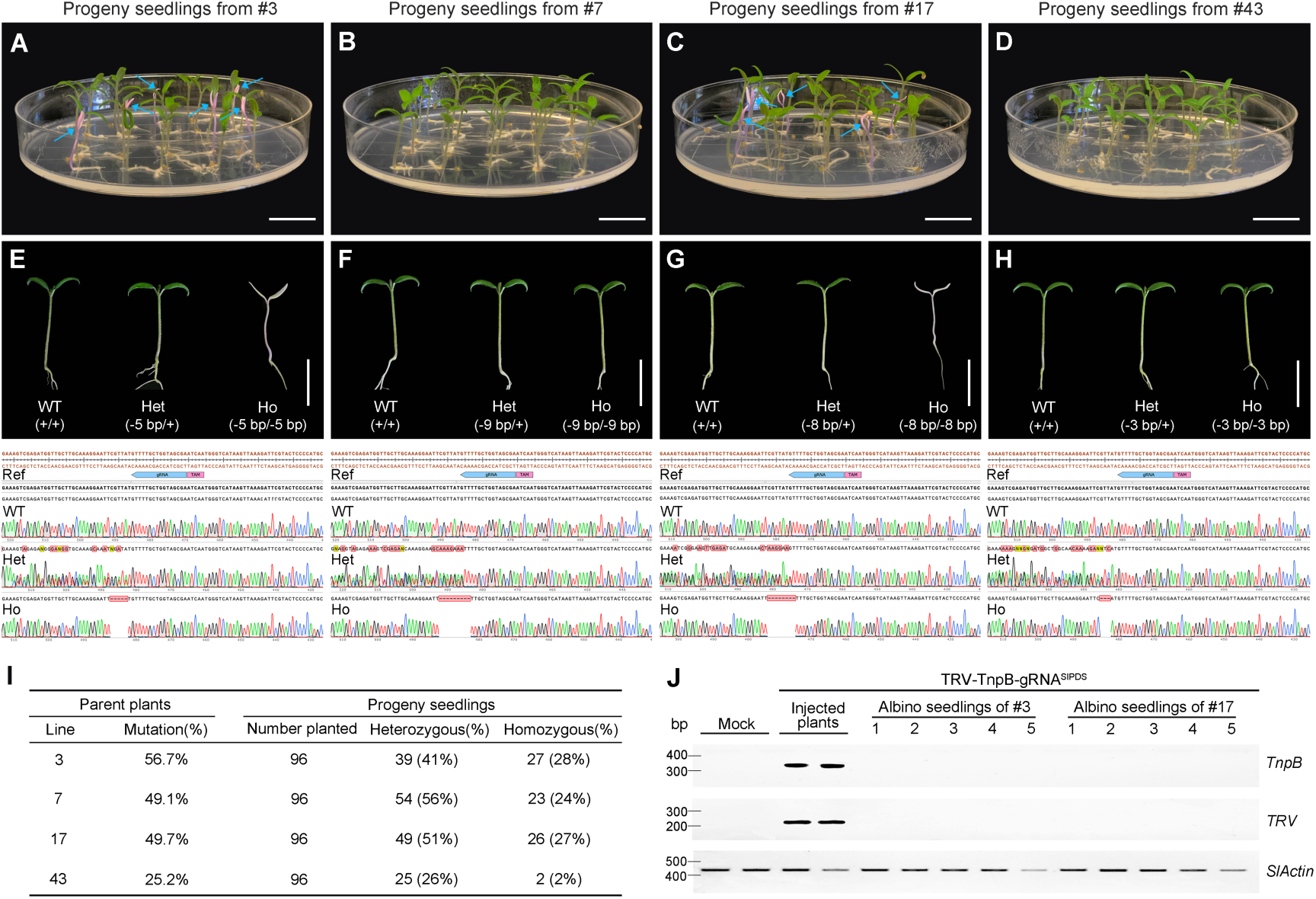
Transgene-free and heritable genome editing in TRV-TnpB-gRNA^SlPDS^-mSlFT infected tomato plants. (**A-D**) Phenotype of 1-week-old progeny seedlings from TRV-TnpB-gRNA^SlPDS^-mSlFT#3 (**A**), #7 (**B**), #17 (**C**) and #43 (**D**) parent plants. The albino seedlings are marked with blue arrows. (Scale bars, 2 cm). (**E-H**) Phenotyping and genotyping analysis of progeny seedlings from TRV-TnpB-gRNA^SlPDS^-mSlFT#3 (**E**), #7 (**F**), #17 (**G**) and #43 (**H**) parent plants. Representative image of 1-week-old progeny seedlings (top panel). Sanger sequencing trace file of PCR product from progeny seedlings (bottom panel). Ref: Reference sequence; WT: wild type; Het: Heterozygous; Ho: Homozygous. (Scale bars: 1 cm). (**I**) Table summarizing the percentage of progeny seedlings with heterozygous and homozygous mutations from the four parent plants shown in panels **A-H**. (**J**) RT-PCR performed using total RNA from two control plants, two TRV injected plants (tissue shortly after injection), and 10 albino progeny seedlings to detect *TnpB* (upper panel) or TRV mRNA (middle panel) expression. The *SlActin* gene served as an endogenous control (bottom panel). The experiment was repeated three times with similar results.

Previous studies have shown that TRV is not transmitted to the next generation in plants(17, 23). To verify the absence of TRV in the progeny of TRV-infected plants, reverse transcription PCR was performed on five albino progenies of both #3 and #17 parent shoots. As expected, TRV and TnpB mRNA signals were not present in any of the albino offspring seedlings (Fig. 2J), and PCR analyses further indicated the absence of viral T-DNA integration (Fig. S8), demonstrating that the TRV-TnpB-gRNA-mSlFT system enables the generation of edited virus- and transgene-free progeny.

### Efficient Somatic and Heritable Editing of *SlDA1* in *De Novo* Shoots Using TRV to Express ISYmu1 and gRNA in Different Tomato Cultivars

Many prior demonstrations of novel genome-editing approaches have relied on targeting genes with readily observable phenotypic readout, such as *PDS* (24–26). To evaluate whether our system can support the editing of other genes, particularly agronomically relevant loci, we targeted the *SlDA1* gene of tomato, whose homologous genes play an important role in the regulation of seed and organ size in other plant species(27–30), but remain uncharacterized in tomato. To this end, we first conducted a BLAST search using the *Arabidopsis* DA1 protein sequence as a query against the tomato genome database and, combined with phylogenetic analysis, identified *Solyc04g079840* as the likely tomato DA1 ortholog (Fig. 3A), hereafter referred to as *SlDA1*. We then generated a TRV–TnpB-gRNA^SlDA1^-mSlFT construct and utilized the same method (Fig. S2) to infect multiple tomato cultivars, including M82, Ailsa Craig and Sweet-100, to test whether our system functions effectively in different tomato cultivars (Fig. 3B). Approximately 5 weeks after injection, 41 *de novo* shoots were initiated from 27 injected M82 plants (Table S1). Three random leaves from each of the 41 *de novo* shoots were collected for genomic DNA extraction and Sanger sequencing analysis. We observed mixed sequencing peaks near the *SlDA1* target site in four of the TRV–TnpB-gRNA^SlDA1^-mSlFT infected shoots #3, #23, #32 and #38 (Fig. S6A), suggesting the presence of mutations. Further amp-seq analysis of these four shoots detected indel frequencies from 63.36% to 99.7% in the *SlDA1* gene. Notably, shoots #3 and #23 contained more than 99% edited reads consisting of roughly 50% of one type of indel and 50% of another, suggesting that these two shoots are biallelic mutants with two different mutations that arose on the two homologous chromosomes very early during shoot development. Similar editing outcomes were also observed in the Ailsa Craig and Sweet-100. In Ailsa Craig, four edited shoots were identified among 47 *de novo* shoots with indel frequencies ranging from 35.76% to 89.62% (Fig. 3C, S6B and S7B). Similarly, in Sweet 100, two edited shoots were recovered out of 24 shoots with editing efficiencies of 84.03% and 99.42%, respectively (Fig. 3C, S6C and S7C).

**Fig. 3.**
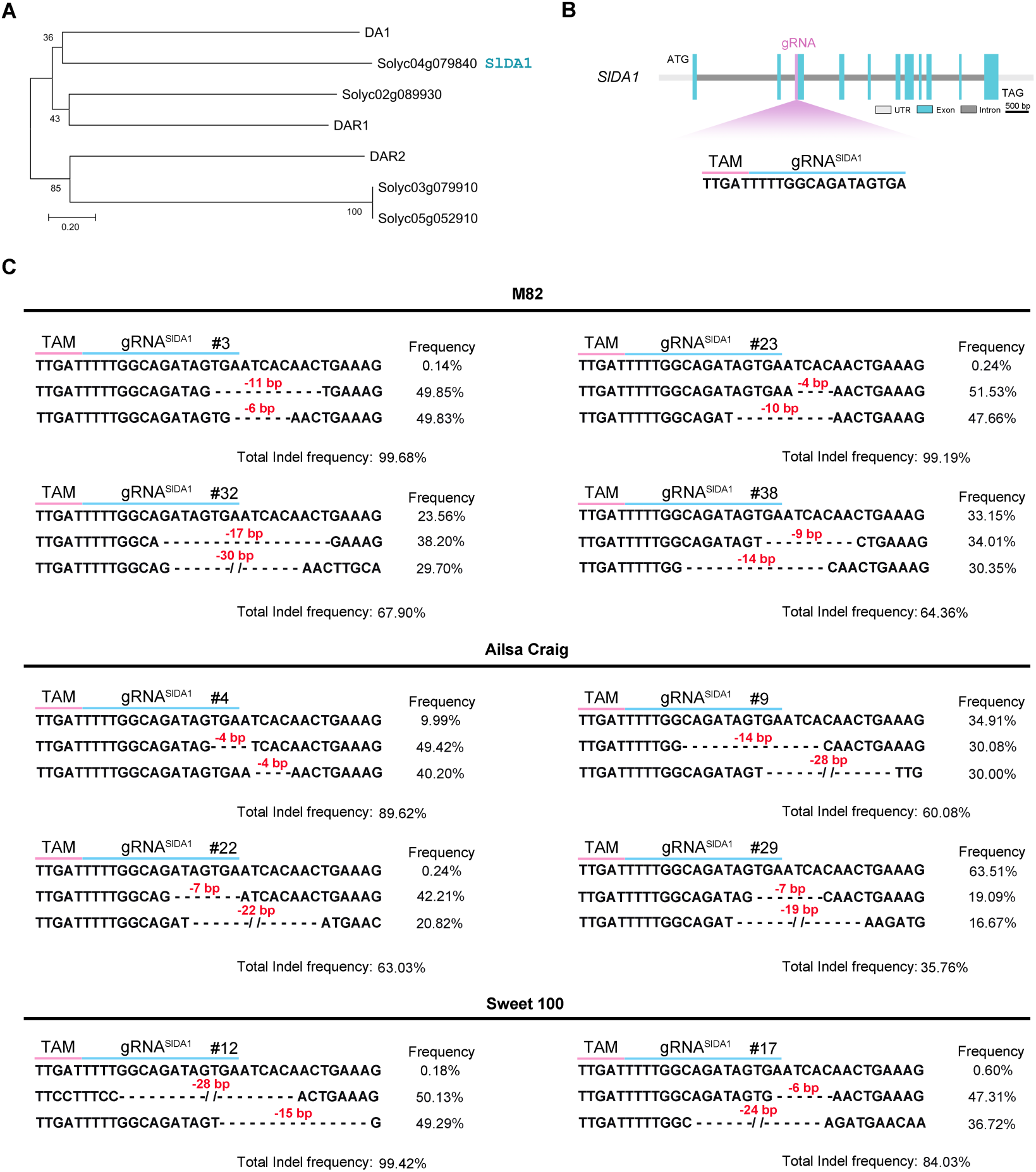
TRV-mediated delivery of TnpB and guide RNA enables somatic editing of *SlDA1* in different tomato cultivars. (**A**) Phylogenetic analysis of tomato and *Arabidopsis* DA1 homologs. SlDA1 is marked in cyan color. (**B**) Schematic diagram of genomic structure. The cyan box represents exons, light grey represents the UTR sequence, dark grey represents introns, the purple box represents the gRNA site. The TAM and target sequences are highlighted by the pink and blue lines, respectively. (**C**) Indel frequency analysis of edited shoots in different tomato cultivars infected with TRV-TnpB-gRNA^SlDA1^-mSlFT. Genomic DNA was extracted from green leaf tissue and subjected to amplicon sequencing using next-generation sequencing (NGS). The TAM and target sequences are highlighted by pink and blue lines, respectively. Deleted residues are marked by black dotted lines. The different indel types are shown on the left. The percentage of reads corresponding to each indel type is shown on the right. The total indel efficiency was calculated by the total number of indel reads divided by total number of reads. Mutant sequences with frequencies >5% are shown as well as the total indel frequency.

To assess whether edited alleles are heritable, we first screened progeny derived from TRV–TnpB-gRNA^SlDA1^-mSlFT #3, #23, #32 and #38 plants in the M82 background. For each line, 48 seeds were germinated on plates and analyzed. As no visible phenotypes were observed in the progeny seedlings, Sanger sequencing was performed to examine the presence of targeted mutations. Notably, all seedlings from #3 and #23 plants harbored biallelic or homozygous mutations. In contrast, progeny from line #32 plant exhibited 38% homozygous and 44% biallelic mutations, whereas those from line #38 plant showed 42% homozygous and 38% biallelic mutations (Fig.4A and 4D). Similarly, progeny from Ailsa Craig and Sweet-100 edited plants also carried heritable *SlDA1* mutations at substantial frequencies (Fig.4B-D). Moreover, RT–PCR analysis indicated that neither TRV2 coat protein RNA nor TnpB transcripts were detected in any of the progeny seedlings (Fig. 4E), suggest they were virus free. PCR analysis further confirmed that the absence of viral vector sequences (Fig. S8), indicating that the progeny did not contain T-DNA insertions. Together, these results demonstrate that TRV-TnpB-gRNA-mSlFT-mediated transgene-free heritable editing can be applied to a gene of agronomic interest and in a variety of different tomato cultivars.

**Fig. 4.**
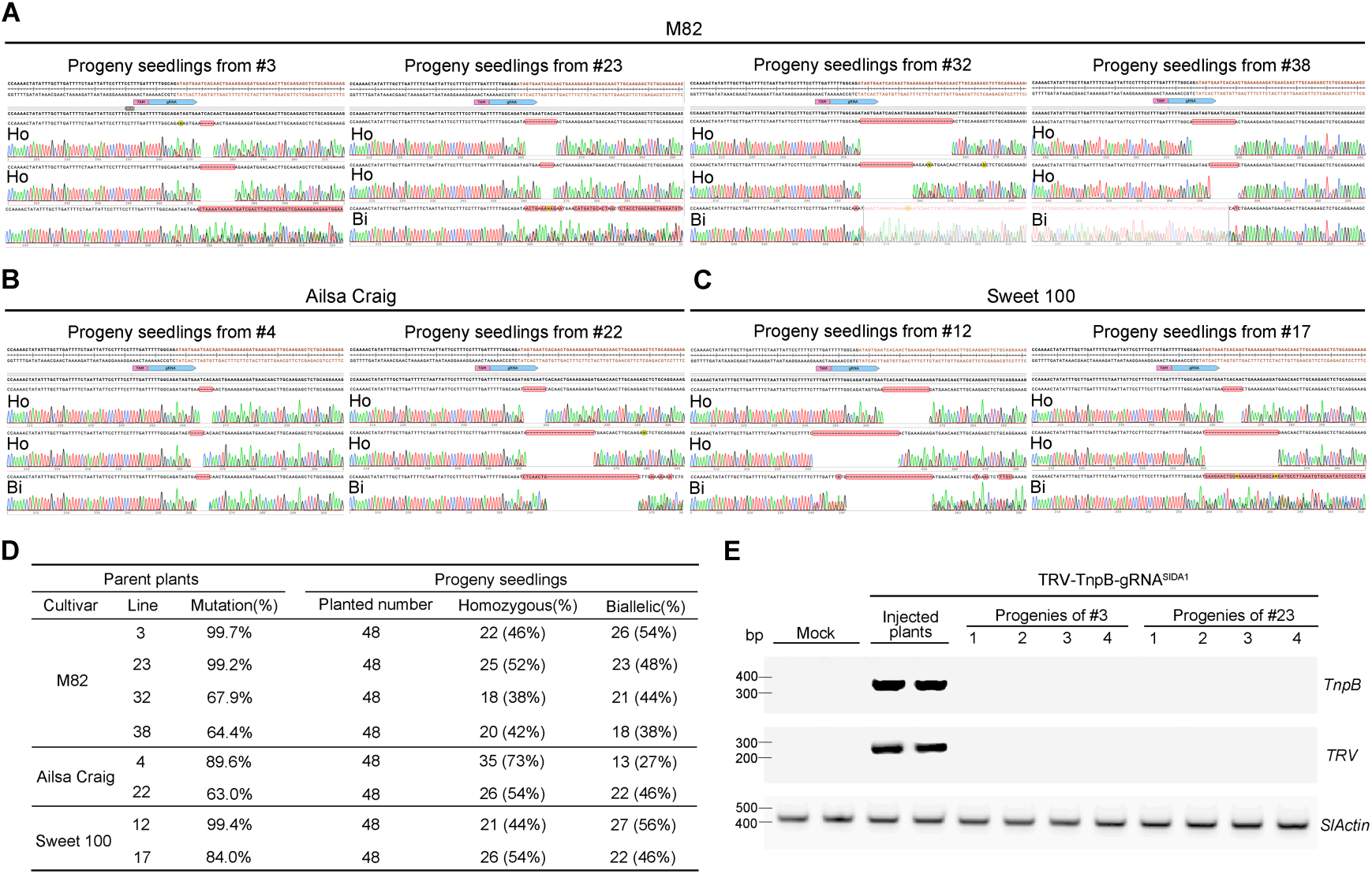
Transgene-free and heritable genome editing in TRV-TnpB-gRNA^SlDA1^-mSlFT infected tomato plants. **(A)** Sanger sequencing trace file of PCR product from progeny seedlings of edited M82 plants. **(B)** Sanger sequencing trace file of PCR product from progeny seedlings of edited Ailsa Craig plants. **(C)** Sanger sequencing trace file of PCR product from progeny seedlings of edited Sweet 100 plants. Ho: Homozygous; Bi: Biallelic. **(D)** Frequencies of biallelic or homozygous mutations in progeny seedlings. **(E)** RT-PCR performed using total RNA from two control plants, two TRV injected plants (tissue shortly after injection), and four progeny seedlings to detect *TnpB* (upper panel) or TRV mRNA (middle panel) expression. The *SlActin* gene served as an endogenous control (bottom panel). The experiment was repeated three times with similar results.

### Loss of *SlDA1* Function Leads to Enlarged Fruit Phenotypes

Fruit size is a major target trait in modern tomato breeding, as it is closely associated with both yield and commercial value(31, 32). To investigate the phenotypic effects of *SlDA1* mutations, three homozygous mutant lines carrying distinct frameshift mutations were selected for analysis. All three independent *SlDA1* mutant lines exhibited visibly enlarged fruits compared to wild-type plants (Fig. 5A). Quantitative measurements further confirmed these observations. Both fruit height and fruit diameter were significantly increased in the mutants (Fig. 5B–C). Compared with wild-type plants, fruit fresh weight in the *SlDA1* mutants was increased by 22%–30% (Fig. 5D). The flower also appeared visibly larger (5A). Together, these results indicate that disruption of *SlDA1* promotes organ growth in tomato, leading to increased fruit size, consistent with the conserved role of DA1 homologs in regulating organ size of plants(27–30).

**Fig. 5.**
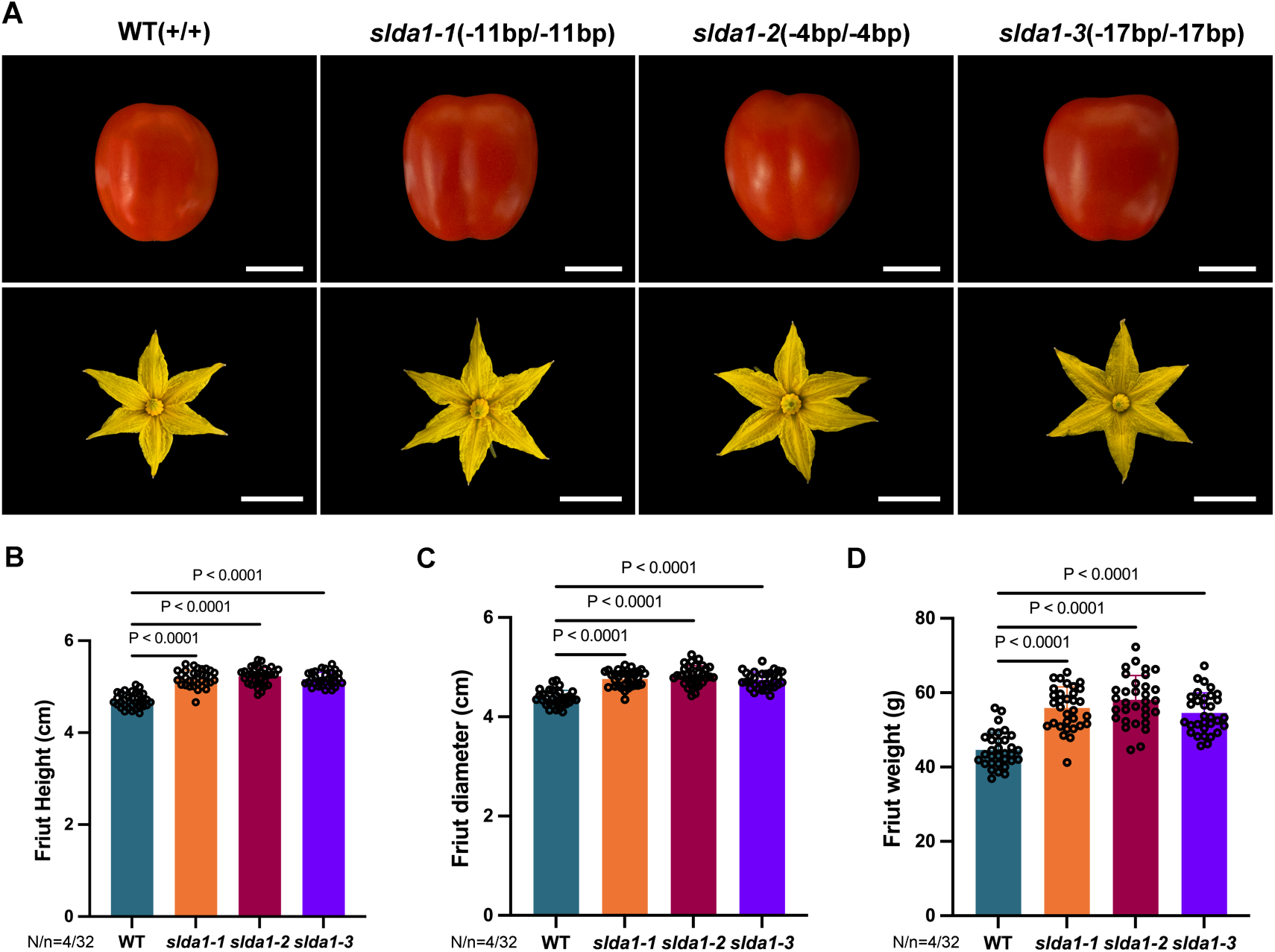
*SlDA1* mutants exhibit increased fruit size in M82 tomato plants. **(A)** Phenotypes of fruits (top panel) and flowers (bottom panel) of WT and different *SlDA1* mutant lines. Scale bars: 2 cm in top panel and 1 cm in middle panel. (B-D) Measurement of height (B), diameter (C) and weight (D) in WT and *SlDA1* mutant plants. Fruits at the red-ripe stage were collected from four independent plants (N=4) per genotype, with eight fruits sampled per plant. Each dot represents an individual fruit, and bars represent the mean ± SE (n=32). Statistical significance compared with WT was determined by one-way ANOVA followed by Dunnett’s test.

## Discussion

To advance modern crop genome editing, it is important to develop rapid and convenient strategies to eliminate the requirement for stable transgene integration and reduce regulatory barriers to generate transgene-free and heritable genome edited crops. VIGE has emerged as a promising approach to achieve these goals(33). In a number of recent studies, tissue culture-free heritable gene editing was achieved by delivering the gRNA to Cas9-expressing tomato plants using TRV(20, 34, 35). However, this approach still requires the generation of Cas9-expressing plants via tissue culture and plant transformation, as well as self- or back-crossing to remove the transgenic components after the edits are obtained. In this study, TRV-mediated delivery of TnpB and its gRNA efficiently generated targeted mutations at the *SlPDS* and *SlDA1* genes in *de novo* shoots, with stable transmitted to the progeny. Thus, our method simultaneously overcomes two major obstacles for tomato genome editing: dependence on tissue culture and the requirement of transgenic components. We targeted both the *SlPDS* gene, a commonly used marker gene with an easily scorable phenotype, and the *SlDA1* gene, which has not yet been studied in tomato and whose disruption led to increased fruit size, and showed that the approach worked robustly in four tomato ecotypes. This suggests that our system will be useful for gene function analysis and crop improvement, and potentially be effective in a wide range of tomato cultivars. Additionally, we obtained plants with enlarged fruits in tomato by knocking out *SlDA1*, which also provides a valuable genetic resource for tomato improvement to increase yield.

FT-based mobile RNA elements have been proposed to enhance long-distance transport of genome-editing reagents to the shoot apical meristem (SAM)(13). However, when tomato seedlings were inoculated with the TRV-TnpB-gRNA^SlPDS^-mSlFT vector via cotyledon injection, low levels of somatic editing were observed, and no edited progenies were recovered. These results are consistent with previous studies in both tomato and pepper, in which TRV-mediated delivery of gRNA^PDS^*-*FT into Cas9-expressing plants resulted in partial photobleaching phenotypes that were absent in later-developing leaves at normal growth temperature and failed to yield heritable mutations(34, 36). This phenomenon has been attributed, at least in part, to activation of plant antiviral immune responses that restrict viral replication and systemic movement, thereby limiting viral accumulation and persistence in apical tissues. For example, a very recent study showed that TRV-mediated delivery of CRISPR/AsCas12f achieved heritable, tissue culture- and transgene-free genome editing in tomato only under conditions of RdRPs knockdown or low-temperature growth(37), highlighting the constraints imposed by host antiviral defenses. Moreover, even if low levels of virus reach the shoot apex, the relatively modest editing efficiency of the compact nuclease TnpB may further constrain successful mutagenesis of germline cells. Finally, as TRV replicates and spreads, it can utilize template switching to recombine out the TnpB and or wRNA cargo sequences, resulting in smaller more efficiently replicating viruses that are incapable of genome editing(19).

To overcome these limitations, we adopted a *de novo* shoot–based editing strategy(34), in which genome editing occurs shortly before the initiation and formation of new shoot meristems to circumvent the need for long-distance viral movement, thereby facilitating stable inheritance of targeted mutations. In contrast to flooding-based inoculation approaches previously employed in *Arabidopsis*, which benefit from the small size and tractability of that model species(10, 17), our strategy is less dependent on whole-plant long distance viral movement and may therefore be readily adaptable to larger plants and diverse crop species. Moreover, the TRV vector in this study possesses a broad host range of more than 400 plant species(38), suggesting this approach described here should work in many additional crops. In addition, ISYmu1 is compact in size and can likely be incorporated into other viral vectors with cargo capacities comparable to TRV, such as potato virus X (PVX) and pea early browning virus (PEBV)(7, 39), further expanding the range of crops that could benefit from this approach. Given that *in planta* viral infection strategies have already been demonstrated with many different viruses and plant species, the potential number of systems that could utilize ISYmu1 is large. Our approach should be especially useful in recalcitrant species that are difficult to genetically transform. In the future, further improving editing efficiency and the target sequence range of ISYmu1 and related TnpB enzymes will be essential to fully realize the potential of this strategy. For example, improved TRV VIGE systems have recently been reported using engineered high-activity TnpBs, and multiplexed editing has also been achieved with this system(40–43).

In summary, by integrating the advantages of a compact RNA-guided nuclease, viral infection, and *de novo* shoot regeneration, this work demonstrates a faster, simpler, and more cost-effective heritable genome editing method that could advance functional genomics and precision breeding in tomato and other crops.

## Methods and Materials

### Plant material and growth conditions

In this study, four tomato genotypes were used for viral infection, including a red cherry type tomato (Amazon.com), M82 (USDA), Ailsa Craig (USDA) and Sweet 100 (Zachary Lippman lab). Seeds were soaked in water at 55°C for 20 minutes, then planted in germination trays and placed in a long-day growth chamber (16-h light and 8-h dark; day 25°C/night 21°C; light intensity: 150 μmol/m^2^/s; 50% humidity) for about 3 weeks.

A different procedure was followed for growing plants used for seed harvesting. Two weeks after germination, small seedlings were transferred from germination trays into large pots. They were grown in a greenhouse with long-day conditions (16-h light and 8-h dark; day 24°C/night 21°C; light intensity:150 μmol/m^2^/s; 50% humidity) for about 3 months.

### Plasmid construction

To generate the TRV2-TnpB-wRNA-mSlFT (pYL001) plasmid, the TnpB-wRNA-*ccdb*-HR-mSlFT sequence with 20 bp overlapping as a gene block was synthesized from Integrated DNA Technologies (IDT). The pMK435 vector was digested by SacI (NEB, R3156S) and PmlI (NEB, R0532S) for 2 hours and the largest fragment was purified using QIAQuick Gel Cleanup kit (Qiagen, 28704). Next, the gene block and digested purified pMK435 plasmid fragment were used to make the pYL001 plasmid using NEBuilder HiFi DNA Assembly (NEB, E2621) according to the manufacturer’s protocol. Finally, the reaction was transformed into the ccdB Survival2 T1R Competent Cells (Invitrogen, A10460).

The pYL001 plasmid was used as a base vector to construct the TRV2 vector with *SlPDS* and *SlDA1* gRNA using the Anneal and Golden Gate strategy. First, two target site oligos were synthesized from Integrated DNA Technologies (IDT) and diluted to 100 μM, and then phosphorylated and annealed. Next, the annealed double-stranded DNA and the pYL001 plasmid were used in a Golden Gate reaction using PaqCI^®^ (NEB, R0745) and T4 DNA Ligase (NEB, M202), followed by the Golden Gate Assembly Protocol. Finally, the reaction was transformed into the 10-beta competent *E. coli* (NEB, C3019). All the plasmids were confirmed using Primordium whole-plasmid sequencing (Plasmidsaurus). The target site primers are listed in Table S2

### Viral infection and *de novo* shoot induction

TRV1 and TRV2 related plasmids were transformed into *Agrobacterium* strain GV3101 electrocompetent cells (Goldbio, CC-207) and grown on LB plates with Kan (50 µg/ml) and RIF (30 µg/ml) for 48 hours at 28°C. A single colony was selected and moved to 2 ml LB liquid media with antibiotics and shaken overnight at 28°C. After confirmation by bacterial PCR, 1 ml of Agrobacterium liquid was transferred into 100 ml of LB liquid media containing KAN (50 µg/ml), RIF (30 µg/ml), 10mM MES and 25 µM acetosyringone and grown overnight in a 28°C incubator. The next day, the Agrobacterium culture was centrifuged for 15 min at 4000 rpm. The supernatant was discarded and the pellet was resuspended in infection buffer containing 10 mM MgCl2, 10 mM 2-(*N*-morpholino) ethanesulfonic acid and 250 µM acetosyringone to an OD_600_ of 0.6. The *Agrobacterium* cells were then incubated at room temperature for 3 hours in darkness. After approximately three hours, the TRV1 and TRV2 Agrobacterium were mixed in a 1:1 ratio and prepared for injection into the tomato plants.

Three-week-old tomato seedlings with about four true leaves were pruned, removing the main shoot and all the axillary shoots. Only leaving two true leaves and two cotyledons. TRV1 and TRV2 *Agrobacterium* mix was injected near the axillary site using 1 ml syringes. Successful injection was confirmed by observing the *Agrobacterium* buffer seeping from the axillary site. Plants were then placed in a dark and 50% humidity growth chamber at 25°C for 2 days, and then grown at 20°C for 2 days. Next, plants were grown in a greenhouse at 16/8-hour day/night cycle at 24/21°C. We removed any new shoots that appeared at the axillary sites in the first month to promote the *de novo* shoot initiation.

### Edited shoots screening and next-generation amplicon sequencing

Approximately 35 DPI, 3 random green leaves from each shoot were harvested for extracting genomic DNA by using Qiagen DNeasy Plant Mini Kit (Qiagen, 69106) according to manufacturer instructions. Target sites were amplified using Q5 DNA polymerase (NEB, M0491s) and PCR products were directly sent for Sanger sequencing. If editing was observed, plants were transferred to larger pots. Otherwise, the unedited shoots were cut at their base to promote new shoot initiation.

Deep next-generation amplicon sequencing was performed for analysis of the mutation frequency of edited plants. DNA libraries were prepared by using a 2-step PCR amplification method. For the first round of PCR, target regions were amplified with primers close to the target site using Phusion™ High-Fidelity DNA Polymerases (Thermo Fisher, F530S) and the reaction was performed under the following cycling conditions: 98 °C for 30s, 98 °C for 30 s, 55°C for 20 s, 72 °C for 30s, 25 cycles (Step2); 72 °C for 5 min. Then, the PCR products were purified by using 1.0× Ampure XP bead purification (Beckman Coulter, A63881). Purified PCR products were then used as the template for the second round PCR by using Illumina indexing primers under the following cycling conditions: 98 °C for 30s, 98 °C for 30s, 60°C for 20s, 72 °C for 30s, 12 cycles (Step2); 72 °C for 5 min. Subsequently, the second-round PCR products were purified and quantified. Finally, equal amounts of the second-round PCR products were mixed and single-end next-generation sequencing on the Illumina NovaSeqX platform was performed. The primers are listed in Table S2.

### Mutation frequency analysis

Amplicon sequencing analysis was conducted as previously described(44). Single-end reads were processed by adapter trimming with Trim Galore using default parameters, and the resulting reads were aligned to the target genomic region with BWA (v0.7.17) employing the BWA-MEM algorithm. The resulting BAM files were sorted, indexed, and subsequently analyzed with the CrispRvariants R package (v1.14.0). For each sample, distinct mutation patterns and their associated read counts were extracted using CrispRvariants. Based on the assessment of control samples, a stringent classification criterion was applied; only reads containing indels of ≥1 bp (insertions or deletions of identical size starting at the same position) with a minimum of 10 supporting reads per sample were considered edited. Single-nucleotide variants were excluded from the analysis.

### Progeny genotyping

Seeds were harvested from the editing parent plants and dried in a 37°C incubator for 3 days. Seeds were sterilized using a 55°C water incubator for 10 mins, 75% ethanol for 1 min, and washed three times with sterile water, 3% NaClO for 7 mins. Seeds were then put in 1/2 MS plates and incubated in darkness at 25°C for 3 days, followed by growth with long-day conditions (16-h light and 8-h dark; day 25°C/night 20°C; light intensity:150 μmol/m^2^/s; 50% humidity) for about 5 days. Once the cotyledons were fully expanded, half of a cotyledon was collected for DNA extraction using the Invitrogen Platinum Direct PCR Universal Master Mix (A44647500) following the manufacturer’s protocol. The extracted DNA was subsequently used as a template for PCR, and the PCR products were subjected to Sanger sequencing. Quantification of indel frequencies was performed with the ICE CRISPR Analysis Tool.

Genotypes were classified according to indel frequencies as follows: Shoots or seedlings with a total indel frequency of 0–10% were considered wild type (WT); samples with a single unique sequence modification at 35–65%, the nonmodified sequence frequency between 35% and 65%, and the total frequency of single unique sequence modification and nonmodified sequence between 85%∼100% was designated as heterozygous (Het); samples with a single unique sequence modification at 85–100% were classified as homozygous (Ho); and samples with two distinct sequence modifications each at 35–65% and the total modification frequency at 85–100% were designated as biallelic (Bi) for two different alleles.

### RT-PCR

Total RNA from progeny seedlings was extracted using Zymo Research Direct-zol RNA MiniPrep kit (R2052). Total RNA was reverse transcribed to synthesize first-strand cDNA using the Invitrogen SuperScript IV VILO Master Mix (11766050). The synthesized cDNA was diluted and then used for PCR with specific primers to detect the TRV virus and TnpB expression by using New England Biolabs Q5 High-Fidelity 2× Master Mix (M0492L) according to manufacturer instructions. The *SlActin* gene was used as the internal control. PCR products were fractionated by 2% agarose gel electrophoresis. The primers are listed in Table S2.

### Fruit phenotypic measurement and statistical analysis

All wild-type and mutant plants were grown on the same bench in a greenhouse under identical conditions (16-h light and 8-h dark; day 25°C/night 21°C; light intensity: 150 μmol/m^2^/s; 50% humidity). To standardize fruit load, the first inflorescence was removed after flowering, and only the four central flowers were retained on each of the second, third, and fourth inflorescences. Fruits were harvested at the red-ripe stage when they had fully developed red coloration. Fruits used for phenotypic measurements were collected from the second to fourth inflorescences. For wild type and each mutant line, four independent plants were analyzed, and eight fruits per plant were selected for measurements of fruit length, diameter, and fresh weight.

Quantitative data are presented as mean ± SD. Statistical significance among different genotypes was evaluated using one-way analysis of variance (ANOVA). Graphs and statistical analyses were performed using GraphPad Prism (V 10.0).

## Acknowledgments

We thank all members of the Jacobsen Lab for their helpful insights and valuable suggestions. We also thank Mahnaz Akhavan for support with Next generation sequencing at the UCLA Broad Stem Cell Research Center BioSequencing Core Facility. We are grateful for the helpful discussion from members of Jennifer A. Doudna’s lab. This work was supported by an NSF Plant Genome Research Program grant (2334027). S.E.J is an Investigator of the Howard Hughes Medical Institute.

## Notes

### Competing Interest Statement

The authors have declared no competing interest.

### Summary of Updates

Section on results updated, Figure3, 4 and 5 updated, Supplemental files updated.

