## Supporting Information for "Virus-induced transgene- and tissue culture-free heritable genome editing in tomato"

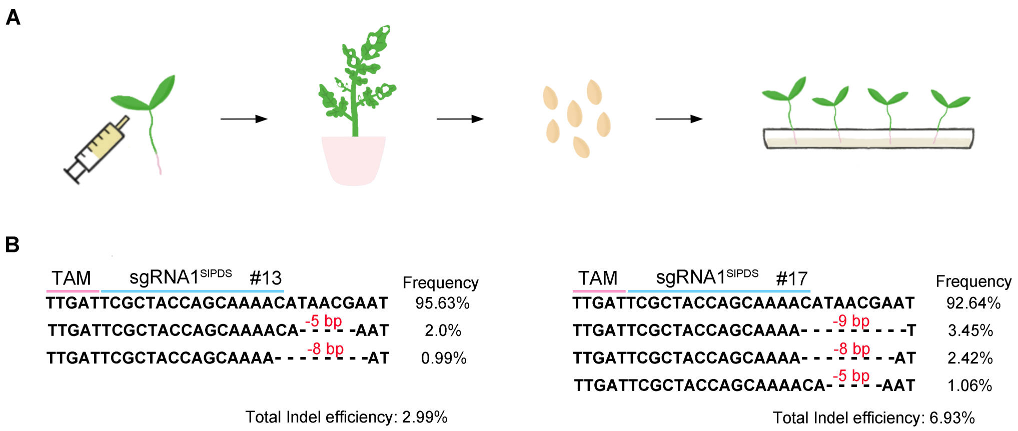


**Fig. S1. Somatic editing in systemically infected leaves of TRV-TnpB-gRNA$^{SlPDS}$ infected tomato plants using cotyledon infiltration. (A)** The procedure of TRV-TnpB-gRNA$^{SlPDS}$-mSlFT mediated genome editing system in tomato plants via *Agrobacterium*-mediated infiltration. *Agrobacterium* cultures harboring both pTRV1 and the TRV-TnpB-gRNA$^{SlPDS}$-mSlFT vectors were co-infiltrated into the cotyledons of 1-week-old wild type plants, followed by sampling of leaves for *SlPDS* edits and collection of seeds for analysis of germline transmission of edits. **(B)** Mutation patterns at the target *SlPDS* site in two TnpB-gRNA$^{SlPDS}$-mSlFT infected tomato plants. Only reads with editing frequency > 0.5% are shown.

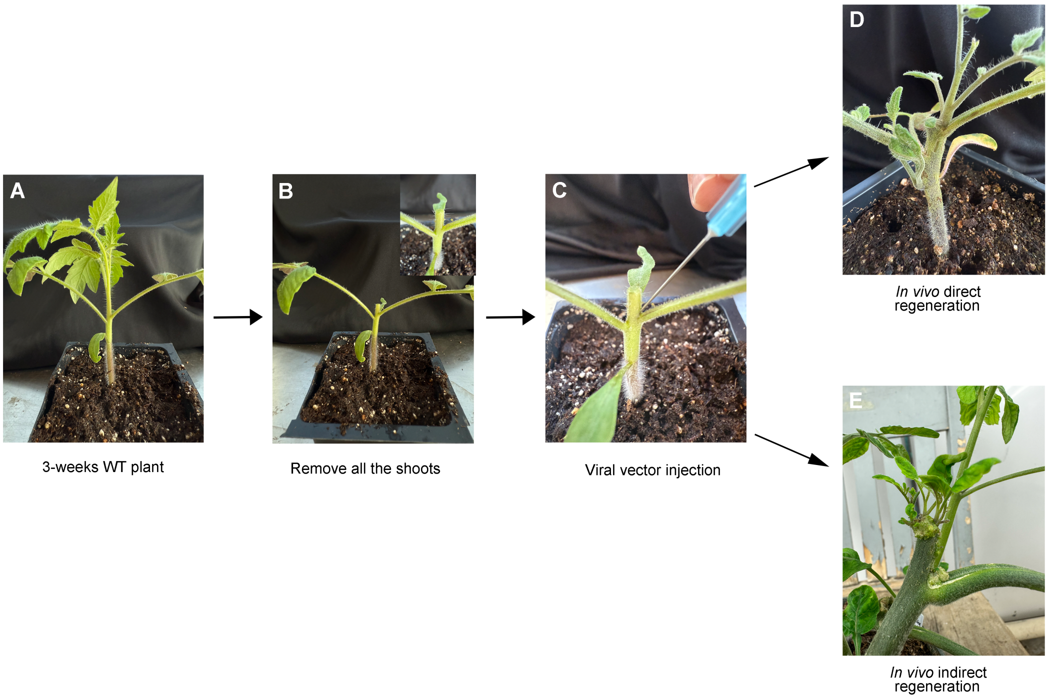


**Fig. S2. Procedure for TRV delivery of ISYmu1 and guide RNA to produce genome editing in *de novo* shoots. (A)** A 3-week-old wild type tomato plant. **(B)** Removal of the top and axillary shoots. **(C)** The viral vector was injected at both the cut stem site and all axils where the axillary meristems had been removed. **(D)** *De novo* shoots directly initiating from axillary regions. **(E)** *De novo* shoots initiating from callus-like tissues.

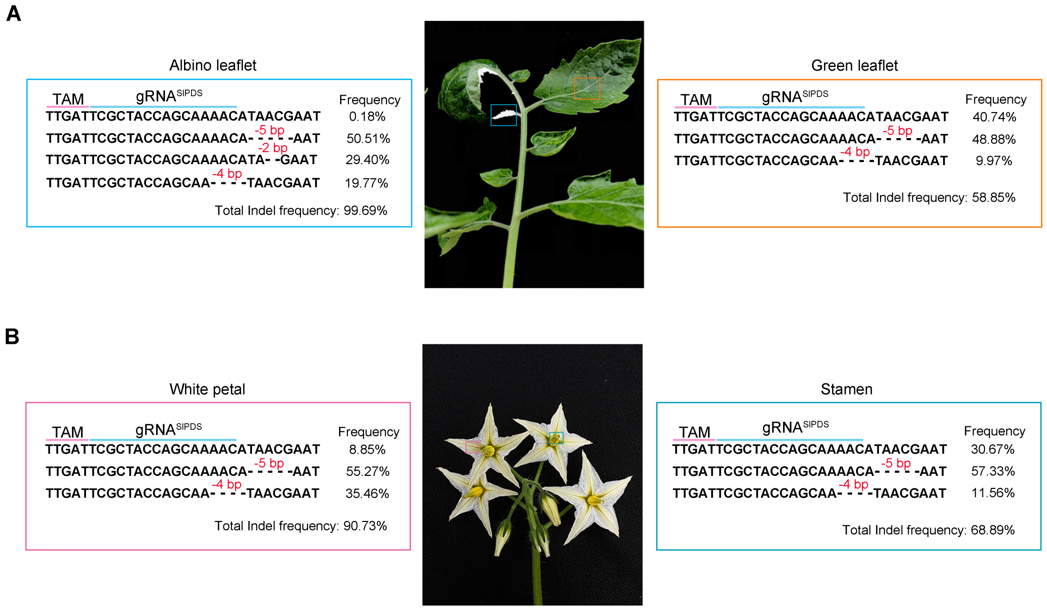


**Fig. S3. PCR-amplicon NGS analysis of different tissues and organs from TRV-TnpB-gRNA$^{SlPDS}$-mSlFT#3 plant. (A)** Mutation patterns and frequencies of green and albino leaflets, respectively. **(B)** Mutation patterns and frequencies of white petals and stamens, respectively. Only reads with editing frequency >5% are shown.

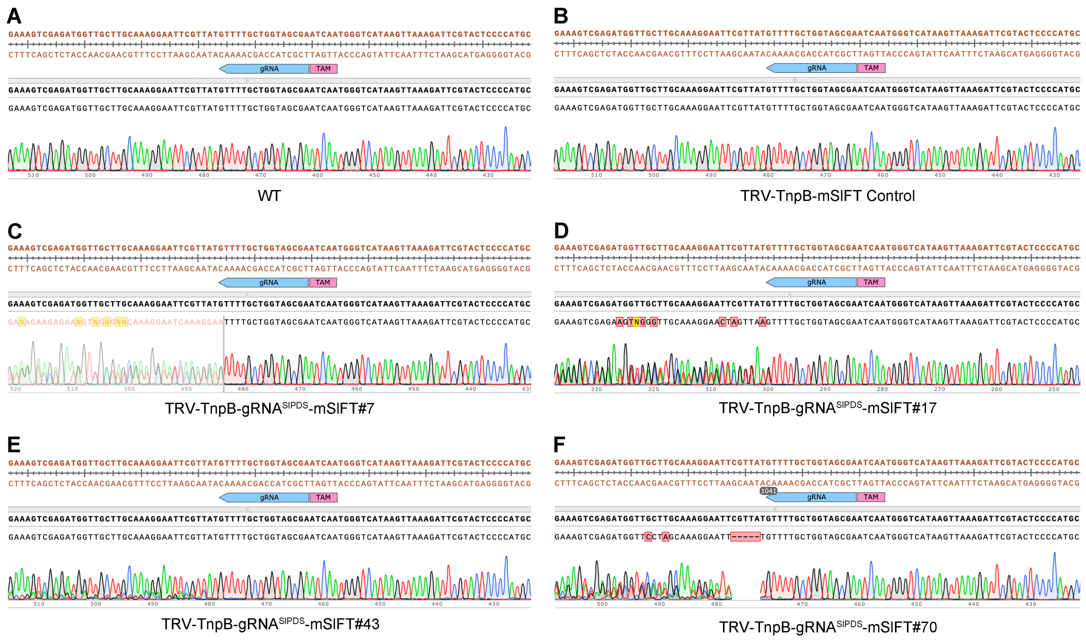


**Fig. S4. Representative Sanger sequencing traces of WT (A), TRV-TnpB-mSlFT Control (B) and five individual edited shoots infected with TRV-TnpB-gRNA$^{SlPDS}$-mSlFT (C-F).** For each panel: Top, reference sequence of *SlPDS* gene; Middle, the sequence of gRNA target and PAM; Bottom, the ab1 trace file. The brown sequence of the reference genome is located in the exon.

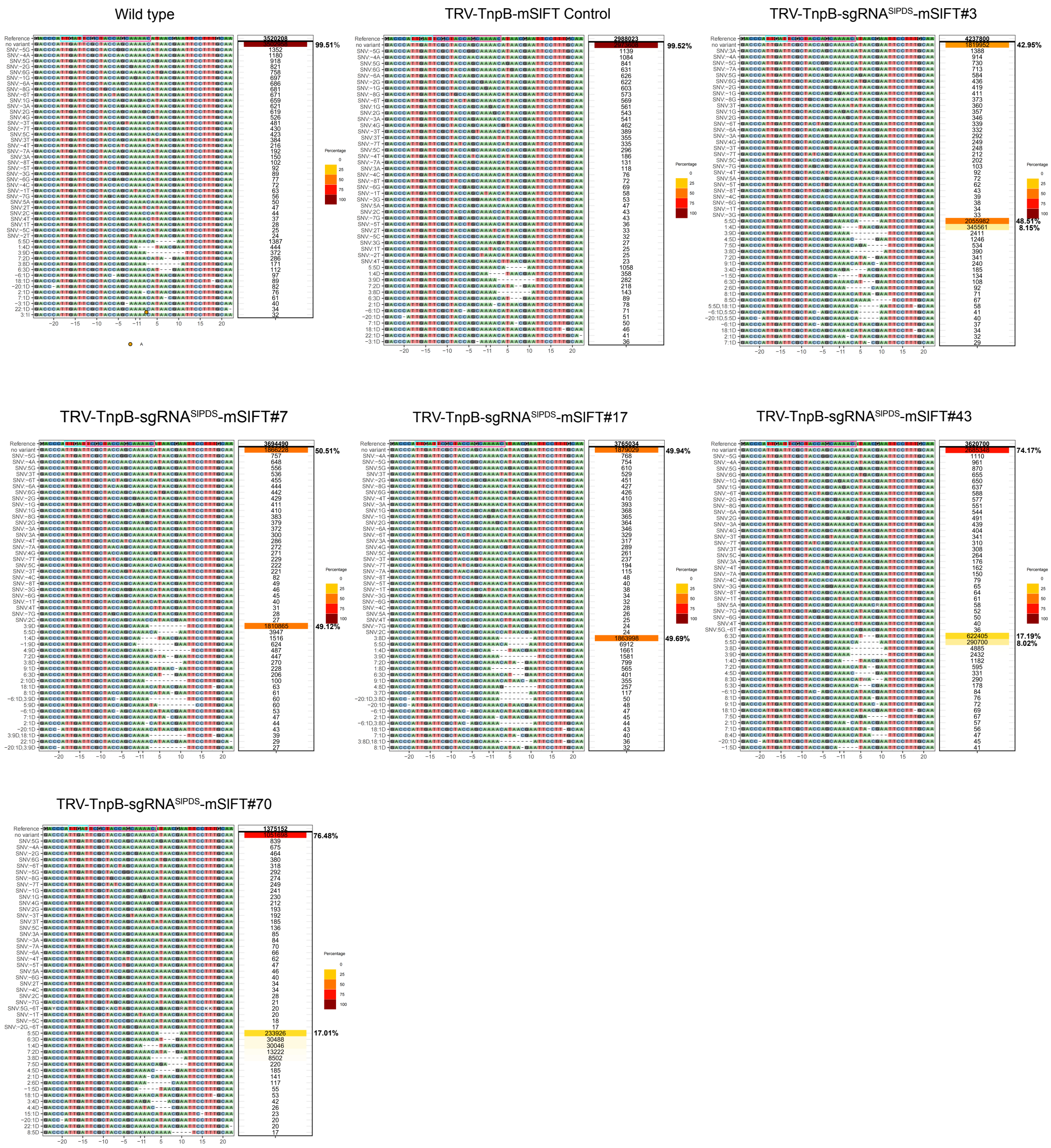


**Fig. S5. PCR-amplicon NGS analysis profile of WT, TRV-TnpB-mSlFT Control and five individual edited shoots infected with TRV-TnpB-gRNA$^{SlPDS}$-mSlFT.** Reads are aligned to the reference sequence, with SNVs and indels annotated on the left. All identified indel variants are included along with their corresponding read counts. Reads for each variant are shown on the right, with the top entry representing the total number of sequencing reads obtained. Indel with editing frequencies >5% are indicated. Color intensity represents the relative abundance (percentage) of each variant.

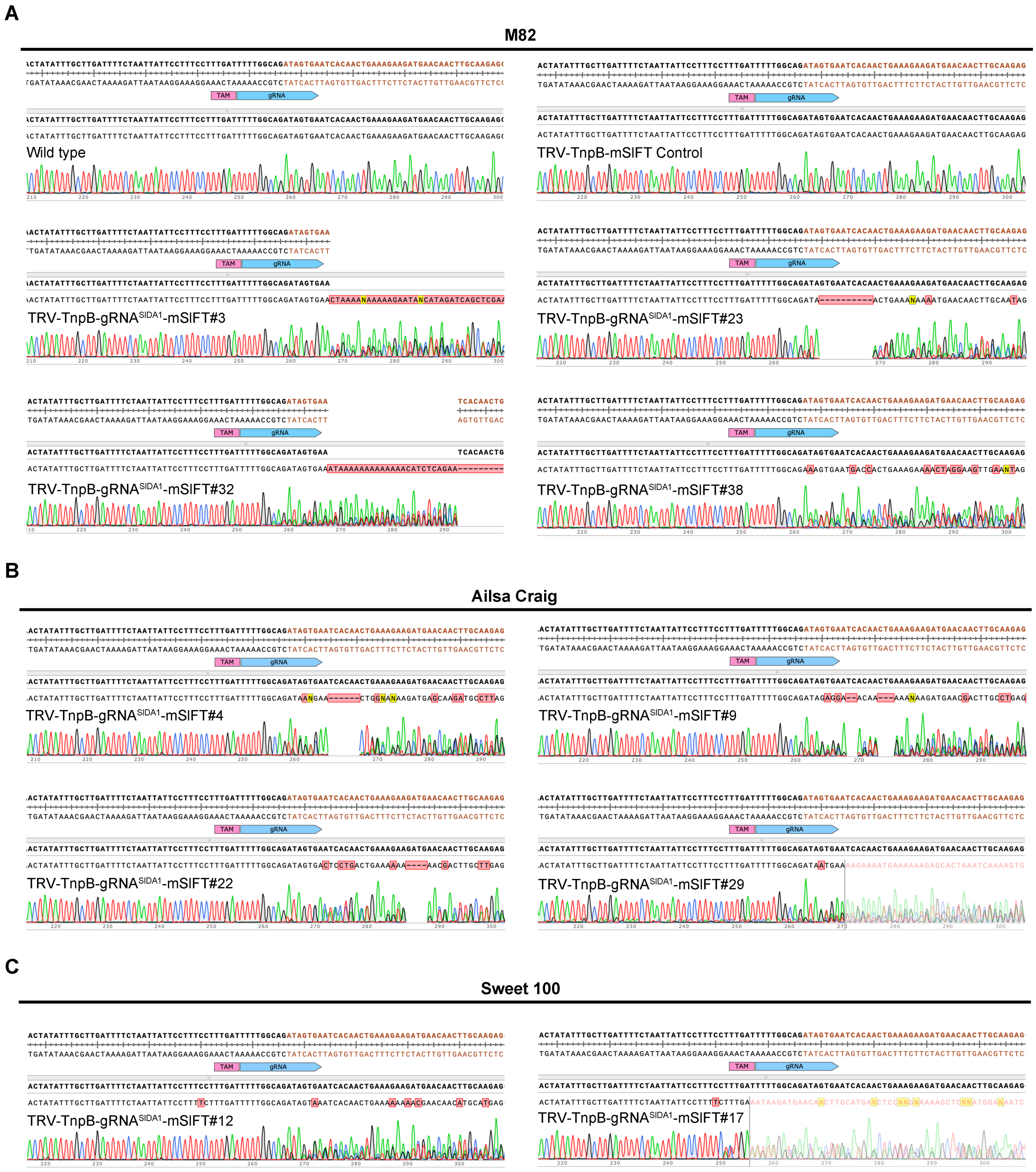


**Fig. S6. Representative Sanger sequencing traces of WT (A), TRV-TnpB-mSlFT Control (A) and individual edited shoots infected with TRV-TnpB-gRNA$^{SlDA1}$-mSlFT in M82 (A), Ailsa Craig (B) and Sweet 100 (C) tomato plants.** For each panel: Top, reference sequence of *SlDA1* gene; Middle, the sequence of gRNA target and PAM; Bottom, the ab1 trace file. The black and brown sequences of the reference genome are located in intron and exon, respectively.

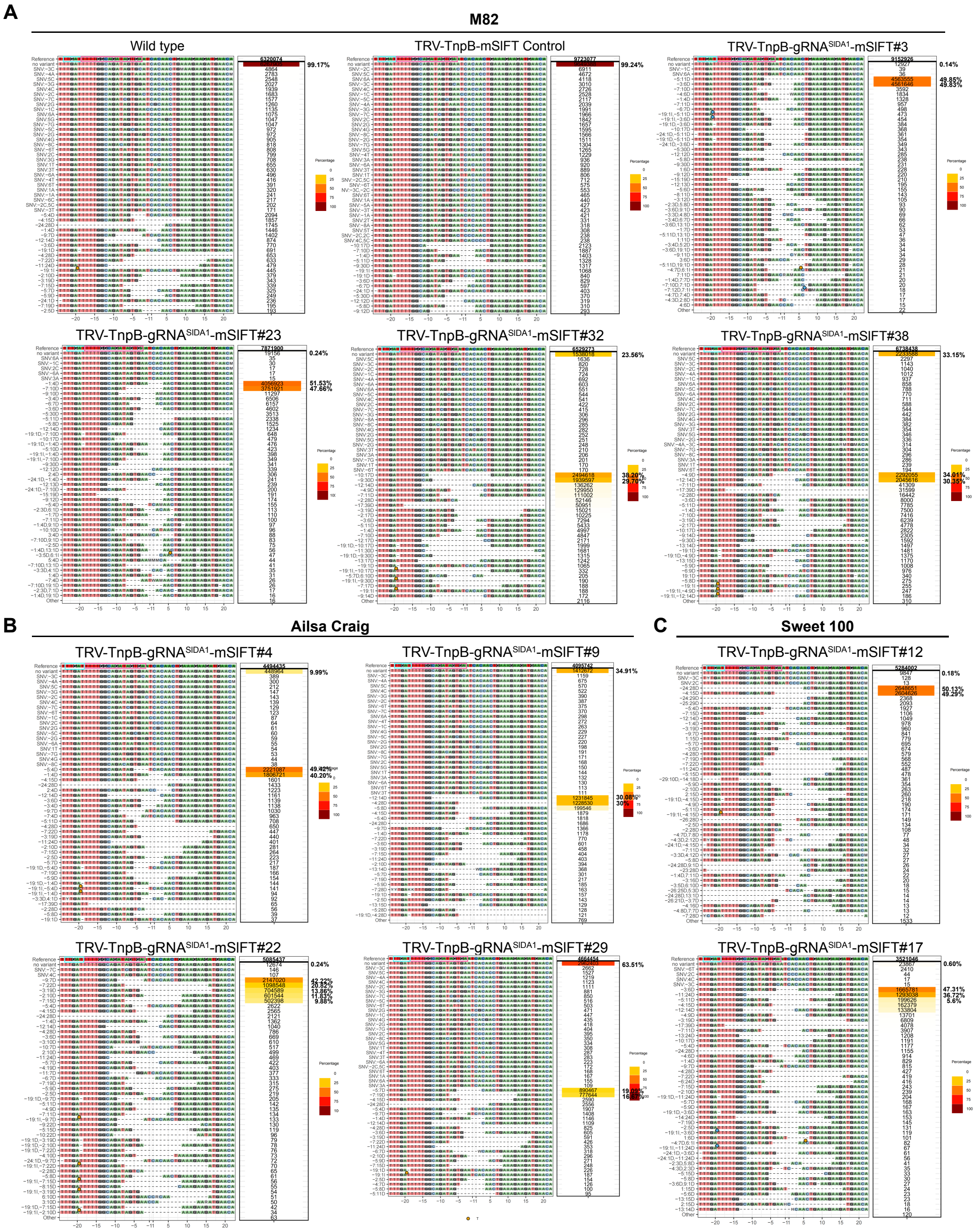


**Fig. S7. PCR-amplicon NGS analysis profile of wild type, TRV-TnpB-mSlFT Control and individual TRV-TnpB-gRNA$^{SlDA1}$-mSlFT infected M82 (A), Ailsa Craig (B), and Sweet 100(C) tomato plants.** Reads are aligned to the reference sequence, with SNVs and indels annotated on the left. All identified indel variants are included along with their corresponding read counts. Reads for each variant are shown on the right, with the top entry representing the total number of sequencing reads obtained. Indel with editing frequencies >5% are indicated. Color intensity represents the relative abundance (percentage) of each variant.

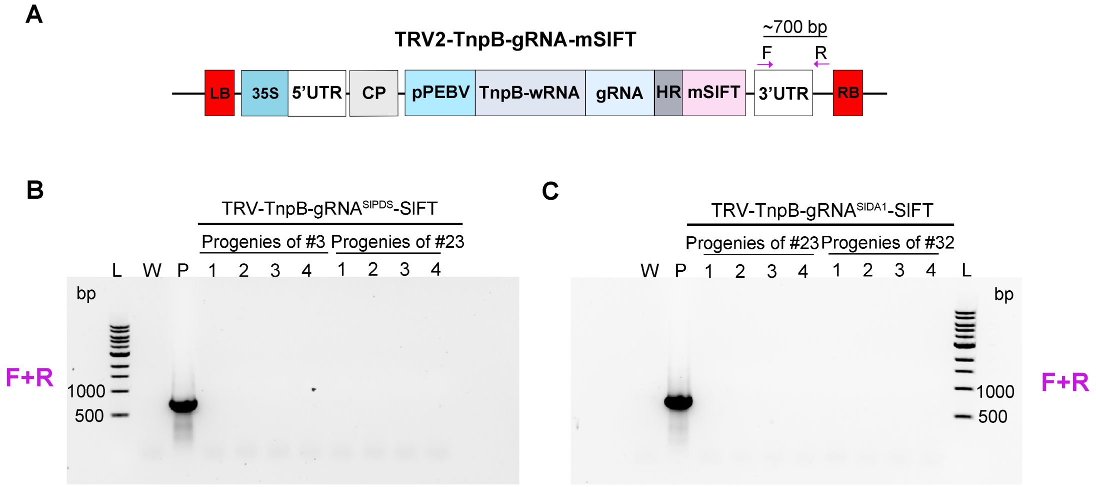


**Fig. S8. Absence of the TRV2-TnpB-gRNA-mSlFT construct in progeny seedlings of edited tomato plants**. **(A)** Schematic diagram of the TRV2-TnpB-gRNA-mSlFT construct. LB: Left Border; RB: Right Border; CP, coat protein; pPEBV, *Pea Early Browning Virus* subgenomic promoter; HR, HDV ribozyme; mSlFT: mutated tomato *FT* mobility element; F: Forward primer; R: Reverse primer. **(B)** PCR gel electrophoresis analysis demonstrating the absence of the TRV2-TnpB-gRNA-mSlFT plasmid sequence in the progeny derived from *SlPDS* edited parent plants #3 and #23 from Red cherry type cultivar. **(C)** PCR gel electrophoresis analysis demonstrating the absence of the TRV2-TnpB-gRNA-mSlFT plasmid sequence in the progeny derived from *SlDA1* edited parent plants (#23 and #32) in the M82 cultivar. L: 1 kb Plus DNA Ladder; P: Plasmid; W: Water; Lanes 1–4: Individual progeny seedlings.

**Table S1. The numbers of injected tomato plants and frequencies of edited shoots**

| Target gene | Cultivar | Injected plants | *De novo* shoots | Edited shoots (%) |
|---|---|---|---|---|
| *SlPDS* | Red cherry-type | 49 | 72 | 5 (6.9%) |
| | M82 | 27 | 47 | 4 (9.8%) |
| *SlDA1* | Ailsa Craig | 31 | 46 | 4 (8.6%) |
| | Sweet 100 | 15 | 24 | 2 (8.3%) |

**Table S2. Primers used in this study**

| **Name** | **Sequences (5'-3')** |
|---|---|
| *SlPDS* gRNA genotyping-F | TCGCAAGTGTGGCTATGGTGGGA |
| *SlPDS* gRNA genotyping-R | CTCCTAGTCCAATCAGCAGTGA |
| *SlDA1* gRNA genotyping-F | AGATGTTCCAGTTGTAGTTGAG |
| *SlDA1* gRNA genotyping-R | ACTATACATATCCACGGCTGAC |
| *SlDA1* NGS-F | ACACTCTTTCCCTACACGACGCTCTTCCGATC TAGTTCAGCTTATCTTTGGAGC |
| *SlDA1* NGS-R | GTGACTGGAGTTCAGACGTGTGCTCTTCCGAT CTTCTGCTTACATGCTGAGGTGG |
| *SlPDS* NGS-F | ACACTCTTTCCCTACACGACGCTCTTCCGATC TAGTTCAGCTTATCTTTGGAGC |
| *SlPDS* NGS-R | GTGACTGGAGTTCAGACGTGTGCTCTTCCGAT CTAGATAATAATTCAAGTCATCAG |
| *SlActin* RT-PCR-F | GGAAAAGCTTGCCTATGTGG |
| *SlActin* RT-PCR-R | CCTGCAGCTTCCATACCAAT |
| *TRV2* RT-PCR-F | TCCTGCTGACTTGATGGACGAT |
| *TRV2* RT-PCR-R | ACCAAGAGCACTGTTAGCCCGTG |
| *TnpB* RT-PCR-F | TGAGGAGACCGGAAAAGGTCTG |
| *TnpB* RT-PCR-R | TCGATTTCCTTCGTACTGTGGCA |
| *TRV2* 3′UTR-F | TGACATTCTCGACTGATCTTG |
| *TRV2* 3′UTR-R | GCGGCACGAAGTCCTATACCAG |
